# Delta-Rhythmic Activity in the Medulla Develops Coincident with Cortical Delta in Sleeping Infant Rats

**DOI:** 10.1101/2023.12.16.572000

**Authors:** Midha Ahmad, Jangjin Kim, Brett Dwyer, Greta Sokoloff, Mark S. Blumberg

**Author notes:** Corresponding Author: Mark Blumberg.

## Abstract

In early development, active sleep is the predominant sleep state before it is supplanted by quiet sleep. In rats, the developmental increase in quiet sleep is accompanied by the sudden emergence of the cortical delta rhythm (0.5-4 Hz) around postnatal day 12 (P12). We sought to explain the emergence of cortical delta by assessing developmental changes in the activity of the parafacial zone (PZ), a medullary structure thought to regulate quiet sleep in adults. We recorded from PZ in P10 and P12 rats and predicted an age-related increase in neural activity during increasing periods of delta-rich cortical activity. Instead, during quiet sleep we discovered sleep-dependent rhythmic spiking activity—with intervening periods of total silence—phase-locked to a local delta rhythm. Moreover, PZ and cortical delta were coherent at P12, but not at P10. PZ delta was also phase-locked to respiration, suggesting sleep-dependent modulation of PZ activity by respiratory pacemakers in the ventral medulla. Disconnecting the main olfactory bulbs from the cortex did not diminish cortical delta, indicating that the influence of respiration on delta at this age is not mediated indirectly through nasal breathing. Finally, we observed an increase in parvalbumin-expressing terminals in PZ across these ages, supporting a role for GABAergic inhibition in PZ’s rhythmicity. The discovery of delta-rhythmic neural activity in the medulla—when cortical delta is also emerging—opens a new path to understanding the brainstem’s role in regulating sleep and synchronizing rhythmic activity throughout the brain.

## Introduction

The brain and body are rich with rhythms.^1–4^ Some rhythms have transparently obvious functions, such as the breathing movements that exchange oxygen and carbon dioxide in the lungs. But the functions of numerous other rhythms, especially brain rhythms, are less obvious. Among these are rhythms that occur during sleep, including the cortical delta rhythm (0.5-4 Hz) that characterizes quiet sleep (QS; or non-REM sleep).^5,6^ Cortical delta is a key player in many aspects of adult sleep. For example, because its amplitude registers the intensity of QS, cortical delta is central to the notion of sleep homeostasis.^7^ Functionally, cortical delta coordinates with other forebrain rhythms to promote learning and neural plasticity.^5,6,8–10^ However, these features of adult sleep are absent in early infancy because cortical delta develops relatively late: In rats, it does not emerge until around postnatal day 12 (P12),^11,12^ at which time QS is just beginning to supersede active sleep (AS; or REM sleep) as the predominant sleep state.^11–14^

For many decades, sleep researchers debated whether the brainstem contains a network for regulating QS that complements networks in the forebrain.^15,16^ Recently, evidence of such brainstem involvement was strengthened by the discovery in adult mice that the parafacial zone (PZ), an area located immediately dorsal to the facial nerve in the medulla, regulates QS.^17,18^ In addition, unit recordings from PZ in adult rats identified state-dependent neurons, including neurons that fire preferentially during QS.^19^ It has been proposed that when GABAergic PZ neurons are activated during QS, they inhibit glutamatergic neurons in the parabrachial nucleus that project to the basal forebrain (BF), resulting in decreased cortical arousal and enabling the expression of cortical delta. ^20^

We hypothesized that cortical delta emerges in part due to developmental changes in PZ and related brainstem circuits that regulate QS. Here, we test this hypothesis by recording PZ and cortical activity in P10 and P12 rats across the sleep-wake cycle. We predicted that developmental changes in the <u>rate</u> of PZ firing would track the onset of cortical delta. Instead, we discovered a striking <u>pattern</u> of PZ activity comprising population-level rhythmic spiking activity during QS. Even more surprising, the rhythmic spiking activity is phase-locked to a local delta rhythm that, in turn, is coherent with cortical delta. The rhythmic activity is also phase-locked with the regular respiration that characterizes QS.^21,22^ These findings introduce an unexpected dimension to the brainstem’s role in sleep regulation and function, raise novel questions about the mechanisms enabling the expression of long-range communication in the infant brain, and open a new path to understanding the intimate connection between a sleep-dependent brain rhythm and the most bodily of rhythms—respiration.^23–26^

## Results

We recorded activity in PZ and frontal cortex using silicon electrodes in unanesthetized, head-fixed P10 and P12 rats as they cycled freely between sleep and wake (**Figure 1A-B**; N = 8 pups/age; P10: 150 units; P12: 136 units; 12-30 units/pup). Analyses were performed only when histology confirmed electrode placements in PZ, as identified previously^20^ (**Figure 1C**). Behavioral states were classified based on respiration, limb movements, and cortical and/or PZ delta (**Figure 1D**; see Methods).

**Figure 1.**
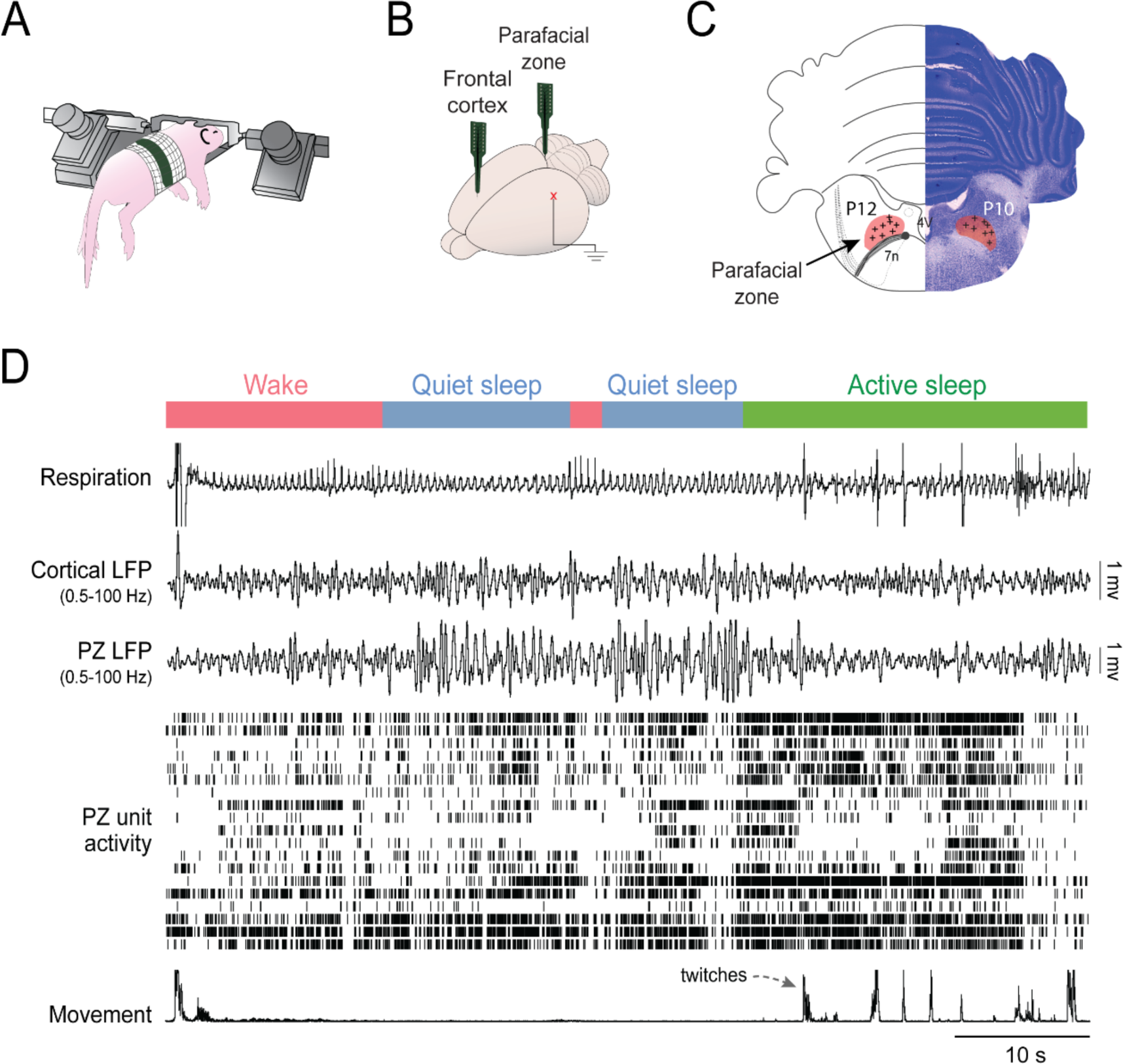
Neural activity in the parafacial zone (PZ) and frontal cortex of infant rats across sleep-wake states. (A) Neural activity and behavior were recorded in head-fixed pups. (B) Silicon electrodes were inserted into PZ and ipsilateral frontal cortex. A ground/reference electrode was inserted into lateral parietal cortex in the contralateral hemisphere. (C) Electrode locations within PZ were confirmed histologically in pups at P12 and P10 (n = 8/age). (D) Representative recording from a P12 rat cycling between wake (red), quiet sleep (blue), and active sleep (green); this color scheme is used for all figures. From top: respiration (inspiration up), cortical and PZ LFP, PZ unit activity, and limb movement. See also **Figure S1A**.

### PZ units exhibit rhythmic spiking during QS at P12

For each PZ unit at P12, we calculated its mean firing rate within each state and then averaged firing rates within pups (**Figure S1A)**. ANOVA revealed a significant effect of behavioral state (*F*_(1.29,9.06)_ = 26.15, *p* < .001, adj. *ηp*^2^ = 0.76), with unexpectedly higher PZ firing rates during AS than during QS or wake. However, closer inspection of activity across all PZ units revealed a striking pattern of rhythmic spiking during QS separated by periods of total silence (**Figure 2A**).

**Figure 2.**
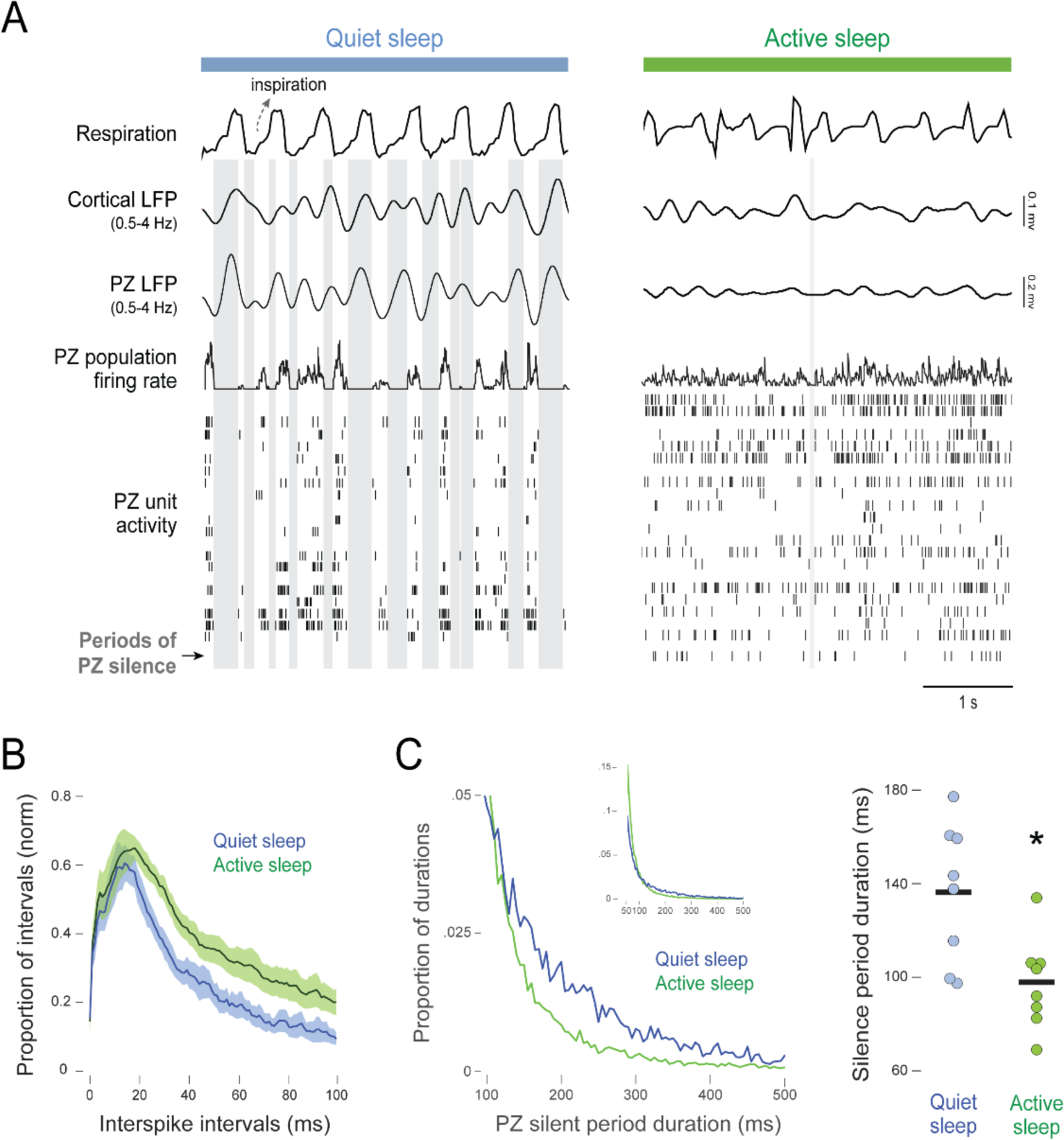
PZ units exhibit rhythmic spiking activity during quiet sleep in P12 rats. (A) Representative 4-s recordings during quiet sleep (left) and active sleep (right). From top: respiration, cortical and PZ LFP, PZ population firing rate (i.e., the mean firing rate across all PZ single units), and PZ unit activity. Gray vertical bars indicate periods in which all PZ units were silent for at least 50 ms. (B) Frequency distribution of mean (± SEM) interspike intervals for PZ single units during quiet and active sleep. Unit data were normalized within state and then averaged within pups. N = 8 pups per plot. (C) Left: Frequency distribution of the duration of PZ silent periods during quiet and active sleep. To highlight state-related differences, only durations between 100 and 500 ms are shown. Data are pooled across all pups. Inset: Frequency distribution of silent periods across the full range of durations from 50 to 500 ms. Right: Mean durations of PZ silent periods during quiet and active sleep. Data points show points for individual pups. Asterisk denotes significant difference. See also **Figures S4 and S5.**

To better understand the nature of PZ’s rhythmic activity, we first compared the frequency distributions of interspike intervals (ISIs) during QS and AS (**Figure 2B**). The higher proportion of longer ISIs (20-100 ms) during AS reflects the higher tonic neural activity during that state. More importantly, the similar ISI peaks (∼20 ms) during AS and QS suggest that PZ rhythmicity during QS does not result from bursts of increased firing. Instead, PZ rhythmicity appears to result from the insertion of regularly occurring periods of total silence. The frequency distribution of these silent periods showed that they were most distinctly associated with QS at durations of approximately 150-400 ms (**Figure 2C, left**). Mean duration of silent periods was significantly longer during QS than AS (*t*_(7)_ = 6.64, *p* < .001, Hedge’s *g* = 2.04; **Figure 2C, right**).

### A local delta rhythm in PZ during QS at P12

In addition to rhythmic spiking, PZ also expresses its own delta rhythm (hereafter, PZ delta). To determine whether PZ and cortical delta have similar properties, we generated mean power spectra in the two structures (**Figure 3A, top**) and quantified delta power for each pup (**Figure 3A, bottom**). Mean delta power was significantly greater during QS than AS in both PZ and cortex (PZ: *t*_(7)_ = 3.65, *p* < .01, Hedge’s *g* = 1.12; cortex: *t*_(6)_ = 9.26, *p* < .001, Hedge’s *g* = 2.95).

**Figure 3.**
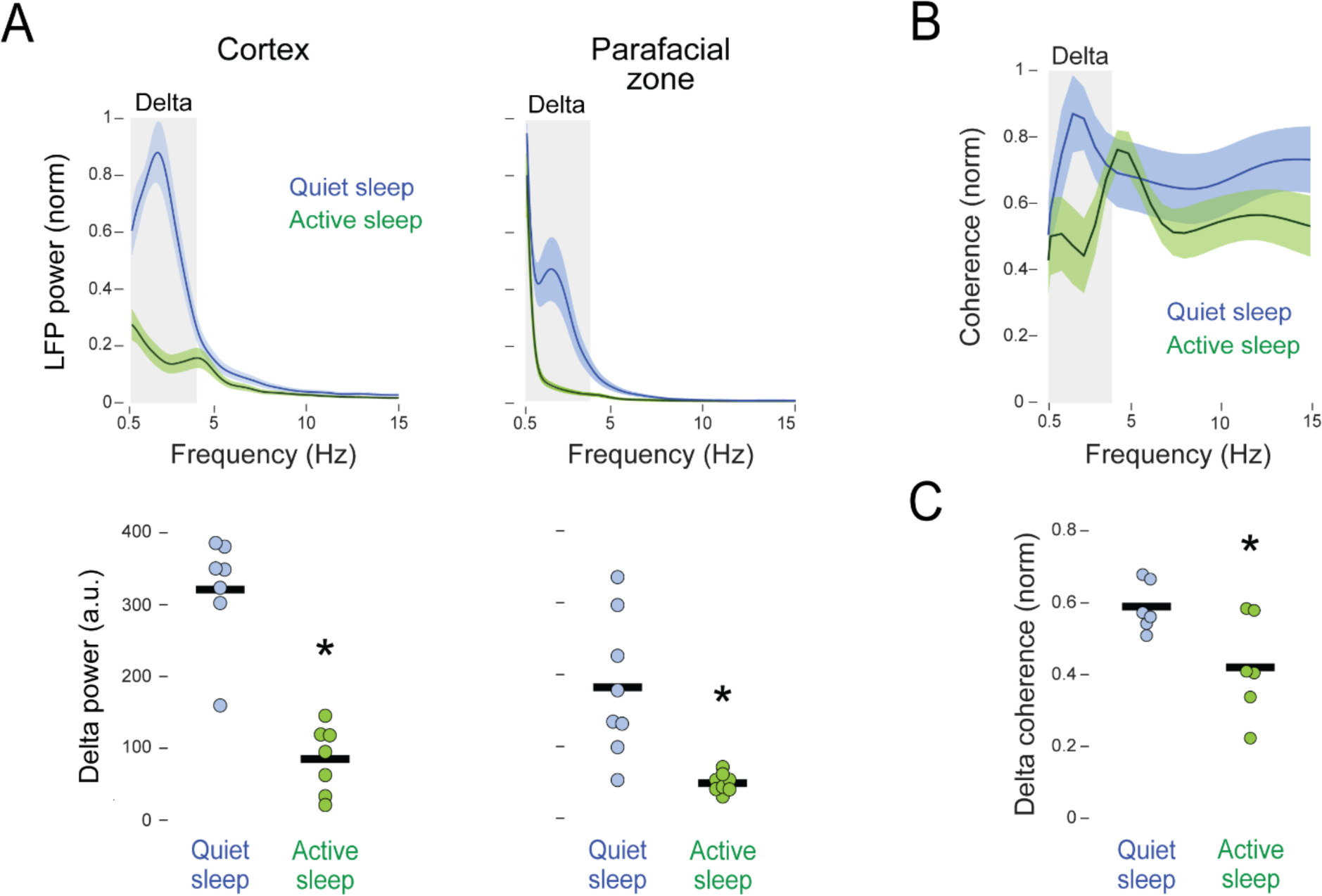
Delta rhythms in PZ and frontal cortex are coherent in P12 rats. (A) Top: Mean (± SEM) LFP power spectra for frontal cortex (left) and PZ (right) during quiet and active sleep. Gray zones indicate delta frequencies (0.5-4 Hz). Data were normalized across states within each structure. Bottom: Mean power (arbitrary units, a.u.) within delta frequencies during quiet and active sleep. Data points show power for individual pups. Asterisks denote significant difference between the two groups. (B) Mean (± SEM) LFP coherence between frontal cortex and PZ during quiet and active sleep. Coherence values were normalized across states. Gray zone indicates delta frequencies (0.5-4 Hz). (C) Mean coherence averaged across delta frequencies in (B), for each pup during quiet and active sleep. Individual data points are average values for pups. Asterisk denotes significant difference between the two groups. See also **Figure S1C**.

The QS-related increase in delta power in PZ and cortex was accompanied by peak coherence at delta frequencies (**Figure 3B**); no such peak in delta coherence was found during AS. Mean delta coherence was significantly greater during QS than AS (t_(5)_ = 5.22*, p* < .01, Hedge’s *g* = 1.70; **Figure 3C**). ^27^

To assess whether PZ delta is produced locally, we computed the distribution of each PZ unit’s activity in relation to the phase of PZ delta. The distributions were averaged within pups before further analysis. During QS, PZ units exhibited a strong tendency to fire near the trough of PZ delta (**Figure 4A, top**). This tendency was not spurious, as no such relation was found during AS (**Figure 4A, bottom**). Phase-locking values (PLVs) of unit activity to PZ delta were computed and z-transformed. During QS, 100% (8/8) of the pups exhibited significant PLVs (*z* > 1.96, *p* < .01); during AS, only 38% (3/8) of the pups exceeded this threshold. We also computed circular concentration coefficients (kappa) of the von Mises distribution (**Figure 4C**); this coefficient reflects the dispersion of the distribution, with higher values indicating tight clustering around specific phases.^27^ Mean kappa values were significantly greater during QS than AS (*t_(7)_* = 7.42*, p* < .001, Hedge’s *g* = 2.28). Thus, we conclude that PZ delta is produced locally.

**Figure 4.**
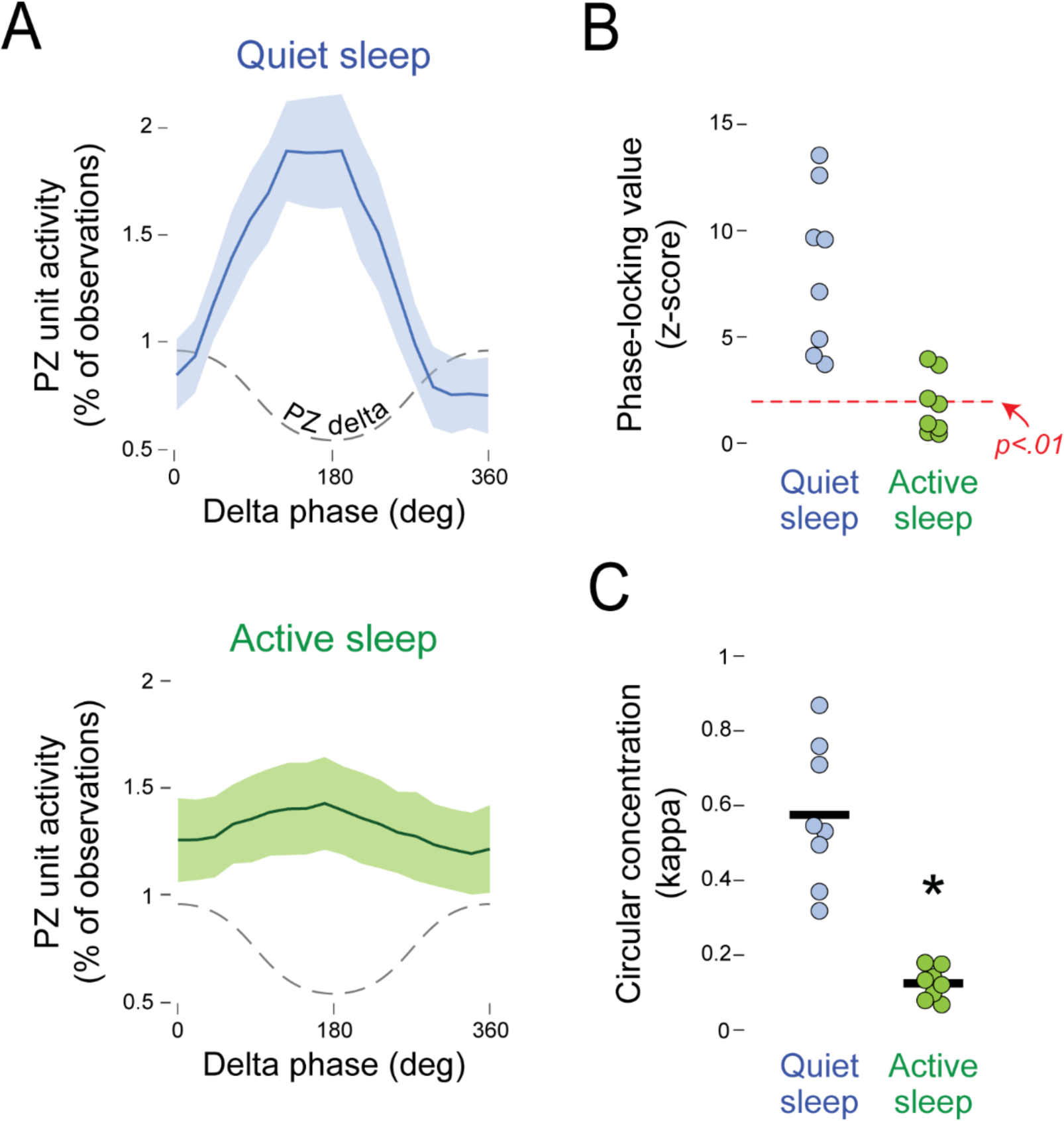
PZ units are phase-locked with local delta during quiet sleep in P12 rats. (A) Mean (± SEM) PZ unit activity in relation to delta phase in PZ during quiet sleep (top) and active sleep (bottom). Delta phase is illustrated with dashed lines. (B) Phase-locking values (z-scored) of PZ units in relation to local delta phase during quiet and active sleep. Individual data points are average values for pups. Red dotted line indicates a significance level of p < .01 based on a z-score of 1.96. (C) Mean circular concentration coefficients (kappa) of PZ unit activity in relation to local delta phase during quiet and active sleep. Individual data points are average values for pups. Asterisk denotes significant difference between the two groups. See also **Figure S2**.

A coherence peak was also found during AS, but at 5 Hz, which is within the range of theta (4-8 Hz) at this age (**Figure 3B**).^28^ However, in contrast with delta, PZ units were not phase-locked with theta (**Figure S2**), suggesting that theta is not generated locally within PZ, but is rather volume-conducted from a nearby region. Phase-locking of unit activity to theta is observed at this age in other structures, including the hippocampus.^28^

### QS-related rhythmic activity in PZ and cortex is weaker at P10 and absent at P8

We also performed recordings at P10, an age when cortical delta is just beginning to emerge and before QS durations begin to increase.^11–14^ To provide a common basis for comparison at P10 and P12, we defined QS as those periods when delta power reached criterion (see Methods). As expected, QS bouts were highly fragmented at P10 in relation to P12 (**Figure S1B**).

Similar to P12, at P10 we saw rhythmic spiking in PZ with intervening periods of silence during QS, accompanied by sporadic bouts of PZ delta and, to a lesser extent, cortical delta (**Figure 5A**). Firing rates at P10 were significantly state dependent (*F*_(2,14)_ = 14.253, *p* < .001, adj. *ηp*^2^ = 0.62) and were again highest during AS, though only significantly higher in relation to wake (**Figure S1A**). The ISI profiles were similar to those at P12 with peaks at ∼20 ms, and evidence of higher tonic firing during AS at longer ISIs (**Figure 5B**); but again, there was no evidence during QS of bursts of increased firing. Silent periods were again more prevalent during QS than AS, but their distribution during QS was more irregular (**Figure 5C, left**) due to fewer silent periods at this age (P10: 8,704 silent periods; P12: 36,678 silent periods). Nonetheless, the mean durations of silent periods were significantly longer during QS than AS (*t*_(7)_ = 8.61, *p* < .001, Hedge’s *g* = 2.65; **Figure 5C, right**).

**Figure 5.**
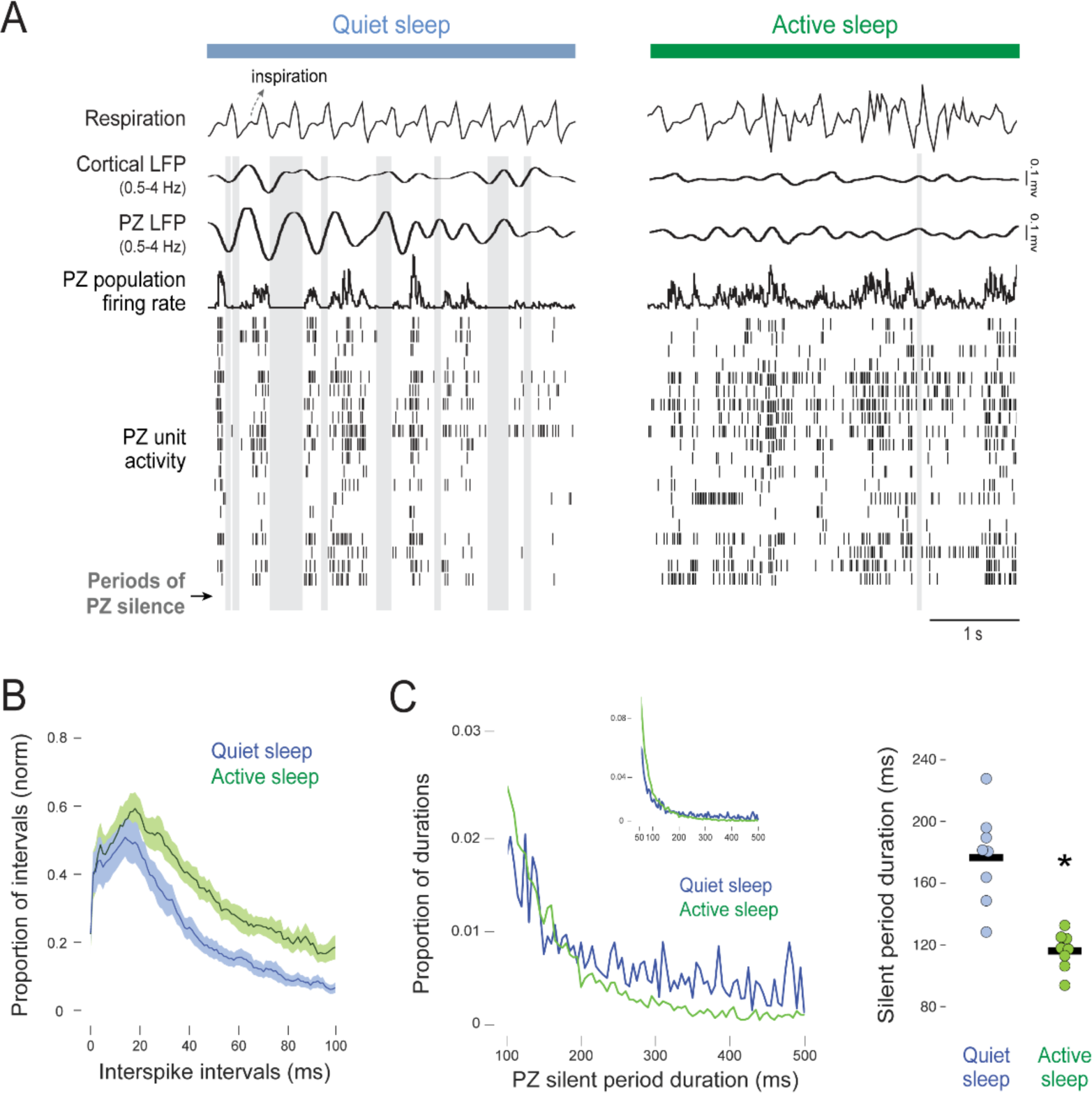
Rhythmic spiking activity in PZ is present but weaker in P10 rats. (A) Representative 4-s recordings during quiet sleep (left) and active sleep (right). From top: respiration, cortical and PZ LFP, PZ population firing rate (i.e., the mean firing rate across all PZ single units), and PZ unit activity. Gray vertical bars indicate periods in which all PZ units were silent for at least 50 ms. (B) Frequency distribution of mean (± SEM) interspike intervals for PZ single units during quiet and active sleep. Unit data were normalized within state and then averaged within pups. N = 7 pups per plot; one pup was excluded due to the absence of single units. (C) Left: Frequency distribution of the duration of PZ silent periods during quiet and active sleep. To highlight state-related differences, only durations between 100 and 500 ms are shown. Data are pooled across all pups. Inset: Frequency distribution of silent periods across the full range of durations from 50 to 500 ms. Right: Mean durations of PZ silent periods during quiet and active sleep. Data points show points for individual pups. Asterisk denotes significant difference. See also **Figures S4 and S5**.

We also compared raw delta power at P10 and P12 (**Figure S1C**). As expected,^11^ cortical delta power increased between P10 and P12. PZ delta power was more pronounced than cortical delta power at P10, and even more so at P12. Thus, at these early ages, PZ may provide a more sensitive indication of QS-related delta than cortex. Further support for this notion comes from the assessment of normalized power spectra at P10 (**Figure 6A**): Mean delta power in PZ was significantly greater during QS than AS (*t_(7)_* = 2.94*, p* < .05, Hedge’s *g* = 0.90), but not in cortex (*t_(7)_* = 1.97, Hedge’s *g* = 0.60). Also, at P10 there was only a weak coherence peak at delta frequencies (**Figure 6B**), and mean delta coherence was not significantly different between QS and AS (*t_(7)_* = 0.145, Hedge’s *g* = 0.04; **Figure 6C**). Finally, PZ units were significantly phase-locked to PZ delta at this age (kappa: *t_(7)_* =2.78*, p* < .05, Hedge’s *g* = 0.86; **Figure S3**).

**Figure 6.**
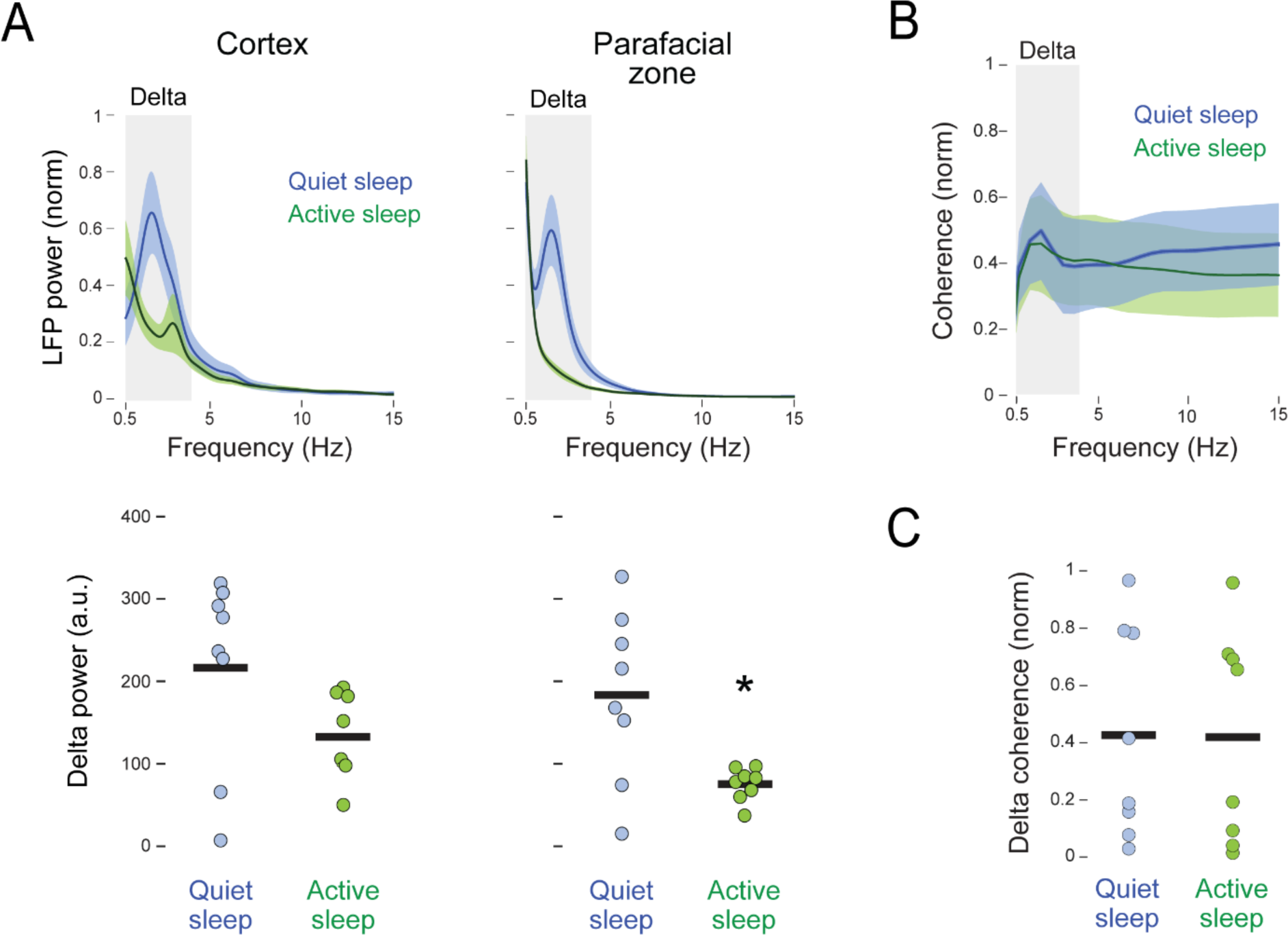
Delta rhythms in cortex and PZ are not coherent in P10 rats. (A) Top: Mean (± SEM) LFP power spectra for frontal cortex (left) and PZ (right) during quiet and active sleep. Gray zones indicate delta frequencies (0.5-4 Hz). Data were normalized across states within each structure. Bottom: Mean power (arbitrary units, a.u.) within delta frequencies during quiet and active sleep. Data points show power for individual pups. Asterisk denotes significant difference between groups. (B) Mean (± SEM) LFP coherence between frontal cortex and PZ during quiet and active sleep. Coherence values were normalized across states. Gray zone indicates delta frequencies (0.5-4 Hz). (C) Mean coherence averaged across delta frequencies in (B), for each pup during quiet and active sleep. Individual data points are average values for pups. See also **Figure S1C**.

The relative strength of PZ delta and the presence of rhythmic spiking activity at P10 leave open the possibility that PZ rhythmicity is already expressed at P8. We tested this possibility by recording PZ and cortical activity in two additional P8 rats (8-10 units/pup). Because cortical delta is not expressed at P8,^11,12^ QS was identified as the period of behavioral quiescence (based on video) between active wake and the first twitch of AS; regular respiration was also used to identify this state. In both pups, we confirmed that there were no clear and sustained periods of PZ or cortical delta, or periods of rhythmic spiking activity (**Figure S4**). Thus, the rhythmic activity in PZ, like that in cortex, begins to emerge at P10. Moreover, the sparse neural activity in PZ at P8 highlights how the emergence of delta-rhythmic activity at later ages depends in part on a general increase in neural activity during QS.

### PZ-related expression of parvalbumin increases between P8 and P12

What factors are responsible for the developmental emergence of PZ delta? Because inhibition is necessary for the expression of many brain rhythms,^29,30^ we hypothesized that inhibitory processing increases in PZ between P8 and P12. This hypothesis derives support from the finding in adult mice that parvalbumin (PV)-expressing inhibitory interneurons are necessary for breathing-entrained rhythmic activity in another medullary structure, the intermediate reticular nucleus (IRt).^31^ In addition, PV-expressing interneurons first appear in the brainstem, cerebellum, and cerebral cortex around P10 and increase rapidly over the next several days.^32–35^ We confirmed that a similar developmental profile of PV expression occurs in PZ between P8 and P12 (n = 3-4 pups/age; **Figure S5**): Whereas PV expression was sparse at P8, we observed increasingly dense labeling of PV terminals at P10 and P12.

### PZ activity is modulated by respiration

Respiratory regularity and irregularity are characteristic of QS and AS, respectively, in infant rats.^21^ We also found here in P12 rats that mean inter-breath interval was significantly longer (*t_(7)_* = 5.50*, p* < .001, Hedge’s *g* = 1.69) and variance was significantly smaller (*t_(7)_* = 12.31*, p* < .001, Hedge’s *g* = 3.79) during QS than AS (**Figure 7A**). Moreover, during QS, respiratory power showed a peak at ∼2 Hz, similar to PZ and cortical delta. During AS, peak respiratory power also peaked within delta frequencies, but the peak was broader and more variable than that during AS, consistent with the irregularity of breathing during this sleep state (**Figure 7B**).

**Figure 7.**
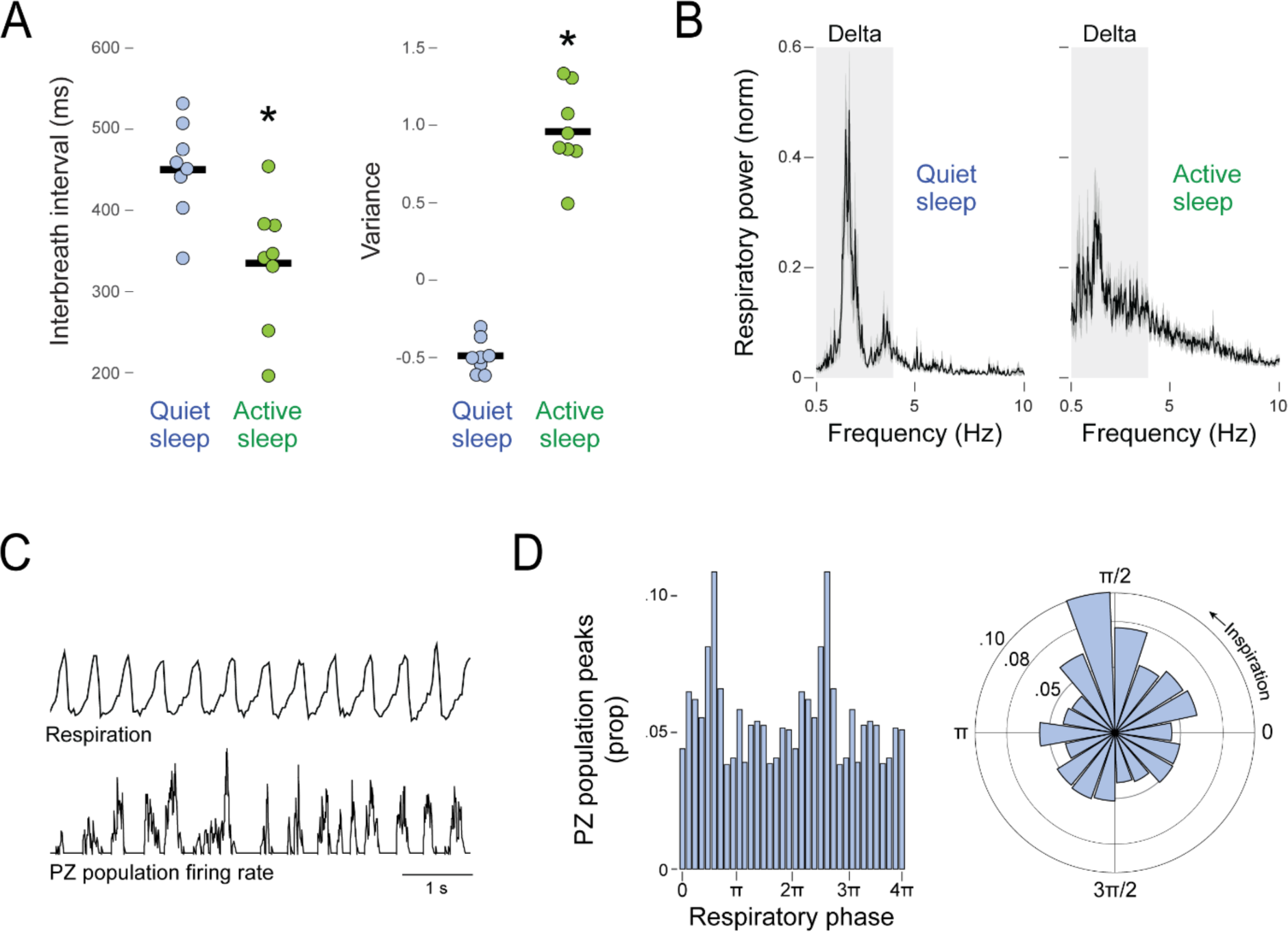
State-dependent modulation of respiration and respiratory modulation of PZ unit activity in P12 rats. (A) Mean inter-breath interval (left) and variance of instantaneous breathing rate (right) during quiet and active sleep. Individual data points are average values for pups. Asterisks denote significant difference between groups. (B) Mean (± SEM) respiratory power spectra during quiet sleep (left) and active sleep (right). Gray zones indicate delta frequencies (0.5-4 Hz). (C) Representative 6-s segment of data during quiet sleep showing a period of regular respiration (top) and associated rhythmic activity in the population of PZ units (bottom). (D) Left: Frequency distribution of mean proportion (prop) of PZ population peaks in relation to respiratory phase during quiet sleep. Note that the data are double plotted. N = 8 pups. Right: Rose plot of the same data at left. The distribution is significantly non-uniform. See also **Figures S6 and S7**.

We next assessed the association between QS-related respiration and PZ population firing rate (**Figure 7C-D**). There were clear and significant peaks in firing rate that occurred after the onset of inspiration (*Z* = 44.43, *p* < .001). This result was obtained using a conservative threshold for defining PZ peak firing rates; similar results were also obtained using more liberal thresholds (**Figure S6**).

### Disconnecting the olfactory bulbs at P12 does not diminish the expression of cortical delta

Respiratory modulation of cortical delta could be mediated directly via ascending projections from PZ (or other brainstem structures) or indirectly via the influence of nasal breathing (**Figure S7A**).^25^ We tested the latter mechanism at P12 by comparing cortical LFPs in an additional cohort of P12 rats with and without bilateral transection of the lateral olfactory tracts (n = 4/group; **Figure S7B-C**). Consistent with a previous study using adult mice,^25^ cortical delta power during QS was not diminished after olfactory transection. That the elimination of olfactory input does not reduce cortical delta power rules out the indirect pathway as a major factor at this age, thereby supporting the notion that a direct pathway is necessary for entraining cortical delta with breathing.^25^

## Discussion

We report in infant rats the unexpected finding of QS-dependent rhythmic spiking activity in PZ, a medullary structure that is implicated in the regulation of QS in adults.^18,20^ PZ’s rhythmic activity is phase-locked to the troughs of a local delta rhythm that, in turn, is coherent with cortical delta. PZ and cortical delta emerge together and become coherent between P10 and P12, suggesting a mechanistic link between the two rhythms. Finally, PZ activity during QS is phase-locked to the regular respiratory rhythm that characterizes that sleep state. Altogether, these findings open the door to a new mechanistic understanding of the brainstem’s contributions to sleep regulation, sleep-related rhythms, and the expression of long-range functional connectivity in early development.

Our findings also inform the decades-long debate about the contributions of the brainstem to cortical rhythmicity.^15^ The delta rhythm is a reliable and prominent feature of forebrain activity during QS,^36^ and this rhythm can be observed in the cortex even after it is surgically disconnected from the brainstem.^16,37^ However, early transection studies also implicated the caudal brainstem in the synchronization of cortical rhythms.^38–40^ The current findings go further than those earlier studies to demonstrate in a medullary structure the presence of rhythmic spiking activity that is phase-locked to a local delta rhythm. Our findings also suggest that the delta-producing system in PZ is mechanistically and functionally associated with the respiratory pacemaker in the ventral medulla.^41^ Thus, we can situate this delta-production system within the broader network of rhythm-generating circuitry in the brainstem, thereby revealing new possibilities for understanding the links among neural oscillations, respiration, and sleep.

### Interpreting PZ’s contributions to sleep and delta

Beginning a decade ago, PZ’s role in the regulation of QS was identified and characterized in adult mice.^17,18^ A subsequent study using adult rats identified PZ neurons with diverse state-dependent profiles, including a plurality (40%) that were selectively active during QS.^19^ However, population-level PZ activity and LFP were not reported in those previous studies, an important difference from the current study that perhaps explains the detection here of rhythmic spiking activity and delta in PZ.

The discovery of rhythmic spiking activity in PZ that is phase-locked with local delta complicates the current model of PZ function described above: That is, the current findings raise the possibility that PZ, beyond its role in regulating QS, also entrains cortical delta. Such entrainment could be mediated via PZ’s ascending direct projections to structures in the hypothalamus, basal forebrain, or thalamus (**Figure S7A**).^42^ Alternatively, perhaps there is no direct causal link between PZ and cortical delta and, instead, the ascending influence of the brainstem on cortical delta arises upstream of PZ. One way to test for PZ’s causal influence is to use optogenetics to briefly perturb its rhythmic activity during QS and assess the immediate effect on cortical delta. Regardless, at this time, the functional significance of PZ delta and the mechanisms that enable its coherence with cortical delta remain to be explained.

### The role of inhibition in PZ’s rhythmic spiking activity

The rhythmic pattern of spiking activity during QS is the most salient feature of PZ activity at P12. Such rhythmicity could result from rhythmic bursting of PZ units and/or rhythmic inhibition. Whereas we did not see evidence of rhythmic bursting, we did see evidence of powerful rhythmic inhibition as reflected in the total absence of spiking activity (as opposed to a relative reduction in activity). Together, these suggest that mean PZ firing rates at P12 would be similar during QS and AS, but for those periods of inhibition during QS that disrupt ongoing activity and reduce the overall mean firing rate (**Figure S1A**). Thus, when interpreting state-dependent differences in firing rates, it is important to also consider the possible confounding effects of state-dependent differences in firing patterns.

The onset of powerful inhibition at P12 may depend in part on two profound changes in inhibitory processing. First, there is the proliferation of PV-expressing inhibitory neurons: Our finding of increased PV expression in PZ between P8 and P12 is consistent with previous findings across the neuraxis.^32–35^ Second, there is the transition at these ages in GABA’s effects on postsynaptic cells from depolarizing to hyperpolarizing.^43–46^ The developmental relations between GABAergic functioning and the expression of delta are not yet established, but a recent study in adult mice suggests that GABA’s postsynaptic effects are dynamically regulated across sleep and wake to influence the expression of cortical delta.^47^

The source of the input that underlies rhythmic spiking activity in PZ is not yet known. A study in adult mice identified strong inputs to GABAergic neurons in PZ from IRt, the deep mesencephalic nucleus (DpMe), the pontine reticular nucleus (PnO), and the lateral hypothalamus (LH).^42^ The last three are notable for their roles in infant^48,49^ and/or adult^50^ sleep regulation, and thus may contribute to the sleep-dependent modulation of PZ. IRt is a large, heterogeneous structure containing neurons that exhibit breathing-entrained rhythmic spiking.^4,31,51^ Thus, we suggest that PZ is part of a larger oscillatory brainstem network that includes IRt and that integrates a diversity of rhythmic behaviors and neural oscillations.^51,52^

### Respiration modulates (sub)cortical delta

The notion that PZ is part of a larger oscillatory network in IRt is further supported by the central role played by respiration.^4^ Evidence for such a role can be traced to the ventral respiratory group, a collection of medullary structures that includes the preBötzinger complex (preBötC), the pacemaker of inspiration.^41^ There exist a diversity of efferent projections from preBötC to brainstem, hypothalamic, and thalamic structures.^53^ There is no known direct projection from preBötC to PZ, but there is one to IRt;^51^ thus, IRt may mediate the respiratory entrainment of PZ rhythmicity.

Considerable effort has been devoted in the past decade to understanding when and how respiration influences the expression of cortical slow waves, including delta, and the functional implications of that influence across sleep-wake states.^23–25,54–56^ Of particular relevance here, it was recently posited that the brainstem conveys respiration-related activity directly to the forebrain, resulting in the entrainment of delta in frontal cortex (**Figure S7A**).^25^ These authors also considered whether respiration entrains cortical delta indirectly via effects on nasal breathing. Because experimental manipulation of activity in the olfactory bulbs reduced but did not prevent respiratory entrainment of cortical delta, it was concluded that entrainment results primarily from a signal, originating in preBötC, that they referred to as a respiratory corollary discharge. In fact, preBötC has few direct projections to the forebrain, but one of its strongest is to mediodorsal thalamus,^53^ which has strong functional ties with frontal cortex.^57^ Similarly, we found here in P12 rats that disconnecting the main olfactory bulbs does not diminish cortical delta power, thus suggesting the predominance of a direct respiratory influence from the brainstem at this age, either directly from preBötC or via other brainstem nuclei such as PZ.

### Conclusions and future directions

Experiments in adult mice using chemogenetic and optogenetic approaches indicated that PZ and nearby structures contribute to the regulation of QS, including delta.^20,58^ At this time, however, we do not know how those manipulations influence rhythmic spiking activity within PZ and the temporal expression of cortical delta. To assess PZ’s functional significance, such data are needed in both infant and adult animals. Future studies should also assess similarities and differences in how PZ and cortical delta develop beyond P12, including their homeostatic responses to sleep deprivation.^59^

Whereas fast brain rhythms (e.g., sleep spindles, gamma) typically reflect local neural processes, slower rhythms are suitable to long-distance communication.^2,29,30^ That delta is expressed coherently between medulla and cortex at P12 is consistent with this general principle governing long-distance communication in the brain. Moreover, the current finding of contemporaneous emergence of delta in PZ and cortex around P12 parallels a similar finding at this age regarding a coherent theta rhythm in hippocampus and red nucleus.^28^ Thus, we now have two examples of slow rhythms that exhibit coordinated activity across forebrain and brainstem, and that emerge during the same brief developmental window. The hierarchical nesting of faster rhythms within slower ones may mirror a similarly hierarchical developmental process, with potential implications for the emergence of behavior and cognition in typically and atypically developing animals.^2,30^ In addition to the emergence of delta and theta rhythms, the P8-to-P12 developmental window includes other major changes in brain and behavior, such as the transition from discontinuous to continuous cortical activity,^60–62^ the proliferation and diversification of inhibitory interneurons,^32,63–65^ an acceleration in the deposition of myelin,^66^ the onset of true whisking^67^, and the locomotor transition to crawling.^68^ Accordingly, the developmental emergence of rhythmic activity in PZ and the onset of delta should be viewed within this broader context of a period of transformational change and, consequently, a period of heightened sensitivity to developmental disruption.

## Acknowledgments

This research was supported by a grant from the National Institutes Health (R37-HD081168) to M.S.B.

## Author Contributions

Conceptualization, M.A., G.S., and M.S.B.; Methodology, M.A., J.K, B.D., G.S., and M.S.B.; Software: M.A. and J.K.; Formal Analysis: M.A. and J.K.; Investigation: M.A. and B.D.; Data Curation: M.A.; Writing – Original Draft, M.A. and M.S.B.; Writing – Review & Editing, M.A., G.S., and M.S.B.; Visualization: M.A., J.K., B.D., G.S., and M.S.B.; Funding Acquisition, G.S. and M.S.B.; Resources, G.S. and M.S.B.; Supervision, G.S. and M.S.B.

## Declaration of Interests

The authors declare no competing interests.

## Inclusion and Diversity

We support inclusive, diverse, and equitable conduct of research.

## STAR METHODS

### RESOURCE AVAILABILITY

#### Lead Contact

Further information and requests for resources should be directed to, and will be fulfilled by, the lead contact, Dr. Mark Blumberg.

#### Materials Availability

This study did not generate new unique reagents.

#### Data and Code Availability

Electrophysiological spiking and LFP data, respiratory data, as well as time codes for states for each animal will be deposited at Dryad for public availability by the date of publication. All original code has been deposited at https://github.com/Blumberg-Lab/Ahmad-et-al.-2024 and is publicly available.

Any additional information required to reanalyze the data reported in this paper is available from the lead contact upon request.

### EXPERIMENTAL MODEL AND SUBJECT DETAILS

Male and female Sprague-Dawley Norway rats *(Rattus norvegicus)* at postnatal day (P) 8, P10, and P12-13 (hereafter P12) were used. Pups were housed and reared in standard laboratory cages (48 x 20 x 26 cm) under 12-h light/dark cycles; food and water were available *ad libitum*. Littermates were always assigned to different experimental groups to avoid litter effects and associated inflation of statistical power.^69,70^ Litters were culled to eight pups by P3. All experiments were performed in compliance with the National Institutes of Health *Guide for the Care and Use of Laboratory Animals* (NIH Publication No. 80–23) and were approved by the Institutional Animal Care and Use Committee at the University of Iowa.

PZ and cortical activity were recorded at P8 (n = 2, 2 males, body weight = 20.1 ± 1.83 g), P10 (n = 8; 5 males; body weight = 23.31 ± 2.47 g) and P12 (n = 8; 4 males; body weight = 29.67 ± 1.41 g). In a separate cohort of pups, PZ and cortical activity were recorded at P12 after bilateral transection of the lateral olfactory tracts (n = 4 pups; 2 males; body weight = 26.25 ± 0.83 g) or sham transection (n = 4 pups; 2 males; body weight = 26.41 ± 0.61 g). Finally, PV expression was assessed in rats at P8 (n = 4; 2 males; body weight = 19.2 + 1.1 g), P10 (n = 3; 2 males; body weight = 22.9 + 1.5 g), and P12 (n = 4; 1 male; body weight = 27.2 + 2.3 g).

### METHOD DETAILS

#### Surgery and Experimental Procedures

##### Neural activity in PZ and cortex across behavioral states

PZ and cortical activity were recorded from unmanipulated rat pups at P8, P10, and P12. As described previously,^71,72^ on the day of testing a pup with a visible milk-band was removed from the litter and anesthetized (3-5% isoflurane gas; Phoenix Pharmaceuticals, Burlingame, CA). The pup was placed on a heating pad for the duration of surgery to maintain body temperature. The scalp was swabbed with iodine and alcohol, and an anti-inflammatory analgesic (Carprofen, 5mg/kg; Putney, Portland, ME) was injected subcutaneously. The scalp was removed and a topical analgesic (bupivacaine, Pfizer, New York, NY) was applied to the skull. Bleach was subsequently applied with an applicator to dry the skull. To prevent bleeding, Vetbond (3M, Minneapolis, MN) was applied around the incision. Two custom-made stainless steel bipolar hook electrodes (50 μm diameter; California Fine Wire, Grover Beach, CA) were inserted into the left and right nuchal muscles and secured with collodion. The pup’s torso was wrapped in soft surgical tape (Micropore, 3M) and, to measure respiration, a piezoelectric sensor (Unimed Electrode Supplies, Surrey, UK) was secured with additional soft tape to the back. Next, the pup’s skull was secured to a stainless-steel head-fix (Neurotar, Helsinki, Finland) using cyanoacrylate adhesive (Loctite, Henkel Corporation, Westlake, OH) and an accelerant (Insta-Set, Bob Smith Industries, Atascadero, CA).

The pup was secured to a stereotaxic apparatus where anesthesia was maintained. Holes in the skull were drilled to enable subsequent electrode placement in PZ (relative to lambda, P10: +1.80 mm rostrocaudal (RC), 1.50 mm mediolateral (ML); P12: +1.80 mm RC, 1.70-1.80 mm ML) and secondary motor cortex (M2; relative to bregma, P10/12: +1.0 mm RC, 1.8–2.0 mm ML). A small hole was drilled over the contralateral parietal cortex for the insertion of a fine-wire thermocouple (Omega Engineering, Stamford, CT) during acclimation and a ground/reference electrode during recording. The ground/reference was a chlorinated silver wire (Ag/AgCl; 0.25 mm diameter, Medwire, Mt. Vernon, NY). Mineral oil was applied to the cortical surface to prevent drying. The duration of the entire surgical procedure was approximately 30 min.

After surgery, the pup was secured to the head-fix apparatus inside a grounded Faraday cage. The pup’s torso was secured using surgical tape to an elevated narrow platform, with limbs dangling freely on each side (**Figure 1A**). Brain temperature was monitored to ensure that it was maintained at 36-37°C; also, the local environment was humidified using moist sponges placed beneath the pup. The pup acclimated to the recording chamber for at least 1 h, and recording did not begin until regular sleep-wake cycles were observed. After acclimation, the thermocouple was replaced with the ground electrode. At that time, data acquisition began, and electrophysiological, respiratory, and video data were recorded for at least 1 h.

##### Cortical delta power during QS after disconnection of the olfactory bulbs from the cortex

PZ and cortical activity were recorded from P12 rats after bilateral transection of the lateral olfactory tracts (n = 4) or sham transection (n = 4), as described previously.^73^ All surgical procedures were identical to those described above, except that the olfactory tracts were transected before the head-fix was attached. Briefly, transections were performed by inserting a 25 G needle through the skull, to the left and right of midline and at the level of the orbits. The needle was inserted perpendicularly until it touched the bottom of the skull and then swept using a side-to-side motion. For sham transections, only the two holes were made. After 90 min of recovery in an incubator maintained at 36°C, the pup was moved to the recording apparatus and electrodes were inserted into PZ and cortex. After at least 1 h and when regular sleep-wake cycles were observed, electrophysiological, respiratory, and video data were recorded for at least 1 h.

#### Electrophysiological Recordings

Neurophysiological and respiratory data were acquired using a data acquisition system (LabRat LR-10; Tucker-Davis Technologies, Alachua, FL). For both recording experiments, PZ data were obtained using a small-site 16-channel linear silicon electrode (Model A1×16-10 mm-100-177-A16; NeuroNexus, Ann Arbor, MI); before insertion, the electrode was coated with fluorescent Dil (Vybrant Dil Cell-Labeling Solution, Invitrogen, Waltham, MA) for subsequent histological verification. PZ electrodes were inserted at a depth of 4.3-4.8 mm. For cortical LFP recordings, a large-site 16-channel linear silicon electrode (Model A1×16-3 mm-100-703-A16; NeuroNexus) was coated with DiI and inserted into M2 at a depth of 0.6–1 mm. Neural signals from the electrodes were sampled at ∼25 kHz using a high-pass filter (0.1 Hz) and a harmonic notch filter (60, 120, and 180 Hz). Respiratory data were sampled at ∼30 Hz.

#### Video Recording and Synchronization

As described previously,^62^ videos of body and limb movements were recorded using a digital video camera (Blackfly-S, 100 frames/s, 3000-ms exposure, 720 x 540 pixel resolution, FLIR Integrated Systems, Wilsonville, OR) and acquired using SpinView software (FLIR). The video was synchronized to the electrophysiological data using an LED visible within the camera frame that automatically pulsed every 3 s for 100 ms. After acquisition, if frames were dropped between LED pulses, a custom MATLAB script was used to insert blank frames.^74^

#### Histology

After recording, the pup was euthanized with ketamine/xylazine (10:1, >0.08 mg/kg IP) and perfused with 0.1 M PBS followed by 4% paraformaldehyde. The brain was extracted and fixed in 4% paraformaldehyde for at least 24 h, after which it was transferred to a 30% sucrose phosphate-buffered solution for another 24-48 h before sectioning. The hindbrain was sectioned coronally at 80 μm using a freezing microtome (Leica Biosystems, Buffalo Grove, IL) and mounted on gelatin-coated slides. Electrode tracks were confirmed using a fluorescent microscope (2.5-5x magnification, Leica Microsystems, Deerfield, IL). Slides were subsequently Nissl-stained using Cresyl violet. Electrode placements were confirmed by overlaying the fluorescent image onto a Nissl-stained image of the same section. For pups with bilateral transection of the lateral olfactory tracts, the brain was extracted from the skull to confirm that the main olfactory bulbs were completely disconnected from the cortex.

#### Pre-Processing of Neural Data

Neural data were preprocessed using custom MATLAB scripts.^62,75,76^ Briefly, to analyze LFPs from PZ and cortex, raw neural activity was downsampled to ∼1000 Hz, smoothed using a .005 s moving Gaussian kernel, and converted to binary files. For unit activity in PZ, raw neural activity was bandpass filtered (300-5000 Hz) and converted to binary files. Putative units were acquired from templates extracted using Kilosort^77^ and visualized and confirmed using Phy2.^78^ Waveforms and waveform autocorrelations were used to identify single units and multi-units (hereafter referred to as units). Sorted units, LFPs, movement data, and respiratory waveforms were imported into Spike2 (Cambridge Electronic Design, Cambridge, UK) or MATLAB for analysis.

#### Classification of Behavioral State

Behavioral state was classified using a combination of respiratory, movement, and LFP data.^11,21,22,74^ As a first step in this process, the respiratory signal was examined for periods with large movement artifacts. Periods where the amplitude of the respiratory signal was 2 standard deviations (SDs) above or below the median were excluded from analysis. To distinguish periods of regular and irregular breathing, respiratory data were analyzed by adapting a previously described method.^22^ Briefly, respiratory peaks were used to derive inter-breath intervals, and these intervals were inverted to obtain the instantaneous breathing rate. Variance of instantaneous breathing rate was computed in 2-s windows and the variances were z-transformed. The 75th percentile was used as a threshold such that values below the threshold identified periods of regular breathing, and values above the threshold identified periods of irregular breathing (consistent with QS and AS, respectively).

Limb movements were detected using a custom MATLAB script that identified changes in pixel intensity between video frames within regions-of-interest (ROIs: forelimbs, hindlimbs, whole-body), producing an output of real-time movement.^74^

Active wake was identified as periods when multiple body parts exhibited coordinated and high-amplitude movements. When a transition to QS occurred, respiration was regular and limb movements were absent. In addition, periods of delta were identified when cortical and PZ LFPs, filtered for delta (0.5-4 Hz), increased above the median LFP amplitude. At P12, both cortical and PZ LFPs were used to define periods of delta; however, because cortical delta was less reliable at P10, only the PZ LFP was used to define periods of delta at this age. AS began with the onset of limb twitching, irregular respiration, and the cessation of cortical and/or PZ delta. If a train of twitches was followed by 10 s of silence, then the AS period ended with the last twitch observed. Finally, periods of wake, QS, and AS were required to be at least 3 s in duration to be analyzed.

#### PV Immunohistochemistry

Brain slices from pups at P8 (n = 4), P10 (n = 3), and P12 (n = 4) were used for PV staining. Brains were sliced in 50 □m sections, with one section used for PV staining and the adjacent section used for Nissl staining. Sections were mounted on gelatin-coated slides and allowed to dry for at least 90 min. Slides were rehydrated in distilled water for 1 min and then permeabilized in 0.2% triton in PBS for 10 min. Slides were washed 3 times for 5 min in PBS, then blocked in 3% BSA blocking buffer (Thermo Scientific, Waltham, MA) with 5% donkey serum (Equitech-Bio, Kervville,TX) for 2 h. Primary antibody against PV (PV27a; Swant, Burgdorf, Switzerland) was diluted 1:100 in blocking buffer and applied to the sections, which were then incubated for 48 h at 4°C. After primary incubation, the slides were washed 3 times in PBS. Secondary antibody (Alexa Fluor 488 donkey anti-rabbit IgG; Life Technologies, Grand Island, NY) was diluted 1:500 in PBS and applied to the sections; sections were then incubated for 90 min at room temperature. The slides were cover-slipped using Prolong Gold (Invitrogen) and expression of PV was visualized and imaged using a fluorescent microscope (Leica).

### QUANTIFICATION AND STATISTICAL ANALYSIS

All statistical analyses were conducted using either SPSS 29 (IBM, Armonk, NY) or MATLAB. Statistical tests included repeated-measures ANOVA, paired t tests, and Rayleigh tests. Data were tested for normality using the Shapiro-Wilks test. When appropriate, Mauchly’s test was used to test for sphericity. In analyses where sphericity was violated, a Huynh-Feldt correction was applied to the degrees of freedom. Bonferroni corrections were used when appropriate to adjust alpha for multiple comparisons. The measure of effect size was partial eta-squared (adjusted for positive bias^79^) for ANOVAs, and as Hedge’s g for paired t tests. Unless otherwise stated, mean data are always presented with their standard error (SEM). Outlier values that were 3 SDs above or below the mean were excluded from analysis; this occurred only once.

#### PZ Firing Rate and Interspike Intervals

To determine if PZ activity was modulated by behavioral state, the average firing rate was calculated for each unit during each state. Unit firing rates were averaged within each pup before statistical analysis.

To generate interspike intervals (ISIs) for single units, a custom MATLAB script was used. Within each state, ISI was normalized to its maximum value. Frequency histograms were then generated using 1-ms bins. Values within each time bin were averaged across pups.

For identifying periods of PZ silence, a custom MATLAB script was used to concatenate unit activity within each pup. Using the concatenated record, silent periods were identified when there was a total absence of activity for at least 50 ms. Normalized frequency histograms of 5-ms bins were then generated to show the distribution of silent periods during each state. Values for each time bin were averaged across pups. The mean duration of the silent periods within each state was then computed.

#### LFP Power Spectra and Coherence in PZ and Cortex

For analysis of LFPs, one electrode channel in PZ and cortex was selected from each pup. Channels were selected based on their location in each region and their raw delta amplitude. In the cortex, higher-amplitude LFPs occurred in the shallower channels.

The MATLAB function, “pspectrum(),” was used to generate raw power spectra. A sampling frequency of 1000 Hz was used, and spectral values between 0.5 and 30 Hz were calculated and normalized within pups across states. The normalized power spectral values were then averaged across pups. One cortical LFP at P12 was excluded due to noise contamination. Finally, the area under the curve within delta frequencies was calculated for each pup to compare delta power between QS and AS.

For generating raw power spectra in olfactory-tract-transected and sham-transected pups, approximately 30-s periods of QS were chosen during which respiration was regular and cortical LFPs (filtered for delta) were above the median amplitude (calculated across the entire recording). These periods were used to generate raw power spectra and delta power, as described above.

For calculating LFP-LFP coherence, we used custom MATLAB scripts.^28,80^ The raw LFPs were bandpass-filtered (0 -.5 Hz) to remove DC components. Each LFP signal was then convolved using a complex Morlet wavelet. To create the Morlet wavelet, the frequency band of interest (0.5-30 Hz) was divided into 50 bins. The temporal resolution of the wavelet was defined using a minimum of 4 cycles and a maximum of 8 cycles. A sampling frequency of 1000 Hz was used, and coherence was generated on a linear scale.

Coherence was computed as follows:

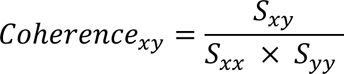

where *S_xy_* is the cross-spectral density between signals *X* and *Y*, and *S_xx_* and *S_yy_* are the auto-spectral densities (i.e., the average power).

Cross-spectral density was computed as follows:

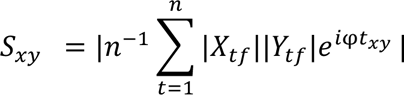

where *S_xy_* contains the complex-valued cross-spectral density. At a specific frequency (*f*) and a given time (*t*), the magnitude of signals *X* and *Y* and their phase differences following Euler’s formula (*e^iφ^*) were computed to obtain the average of magnitude-modulated phase values. For auto-spectral density, all the computations were the same, except the same LFPs were used twice.

Coherence values between cortical and PZ delta (0.5-4 Hz), as calculated above, were normalized within each pup across states, and then averaged across pups. One P12 pup was excluded from the analysis due to a noisy cortical LFP, and another was excluded as an outlier.

#### Phase Locking of PZ Unit Activity with Local Delta

Phase-locking values (PLVs) for PZ units were computed using custom MATLAB scripts.^28,81,82^ Briefly, the phases of the bandpass-filtered (0.5-4 Hz) LFPs were extracted using Hilbert transform. Spike trains for each unit were used to generate spike-phase distributions (phase bin size = 20°); for each LFP phase, the occurrences of spikes were counted and then divided by the total number of spikes.

Rayleigh tests were used to determine significant PLVs.^81^ First, phase estimates were assigned to each index of spike times, and a phase histogram was extracted for each spike. PLVs were computed and z-transformed by bootstrap shuffling (n = 1000 iterations). To do this, a random LFP point was selected and segments on either side of that point were shuffled.

The circular concentration coefficient (kappa; κ) was calculated using the von Mises probability distribution.^27^ The probability density function under this distribution was calculated as follows:

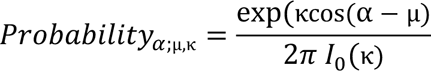

where α is the angular direction, *μ* is a measure of location (i.e., clustering around *μ*), and *I_0_* is the modified Bessel function of order zero. A higher κ indicates that the PLVs were more likely to be concentrated at specific phases of delta.

#### Respiratory Modulation of PZ Unit Activity

Inter-breath intervals and variance of instantaneous breathing rate were calculated as described above for each P12 rat during QS and AS. The spectral power of the respiratory rhythm was generated as described for LFP, normalized across state, and then averaged across pups.

For capturing the respiratory modulation of PZ population firing rate at P12, the average firing rate across all PZ units within a pup was calculated using 1-ms bins. Next, the average PZ firing rate was smoothed using a 5-ms moving window. The values exceeding a criterion were used to generate PLVs of the PZ population firing rate in relation to the respiratory rhythm. Three increasingly conservative criteria were used: 0 x median (no threshold), .25 x median, and .50 x median. PLVs were generated as described above for LFPs, but here the PZ population firing rate was used instead of individual PZ units. The population PZ peaks were mapped along the oscillatory respiratory phases (20°/bin) to generate histograms and rose plots.

---

**Figure S1.**
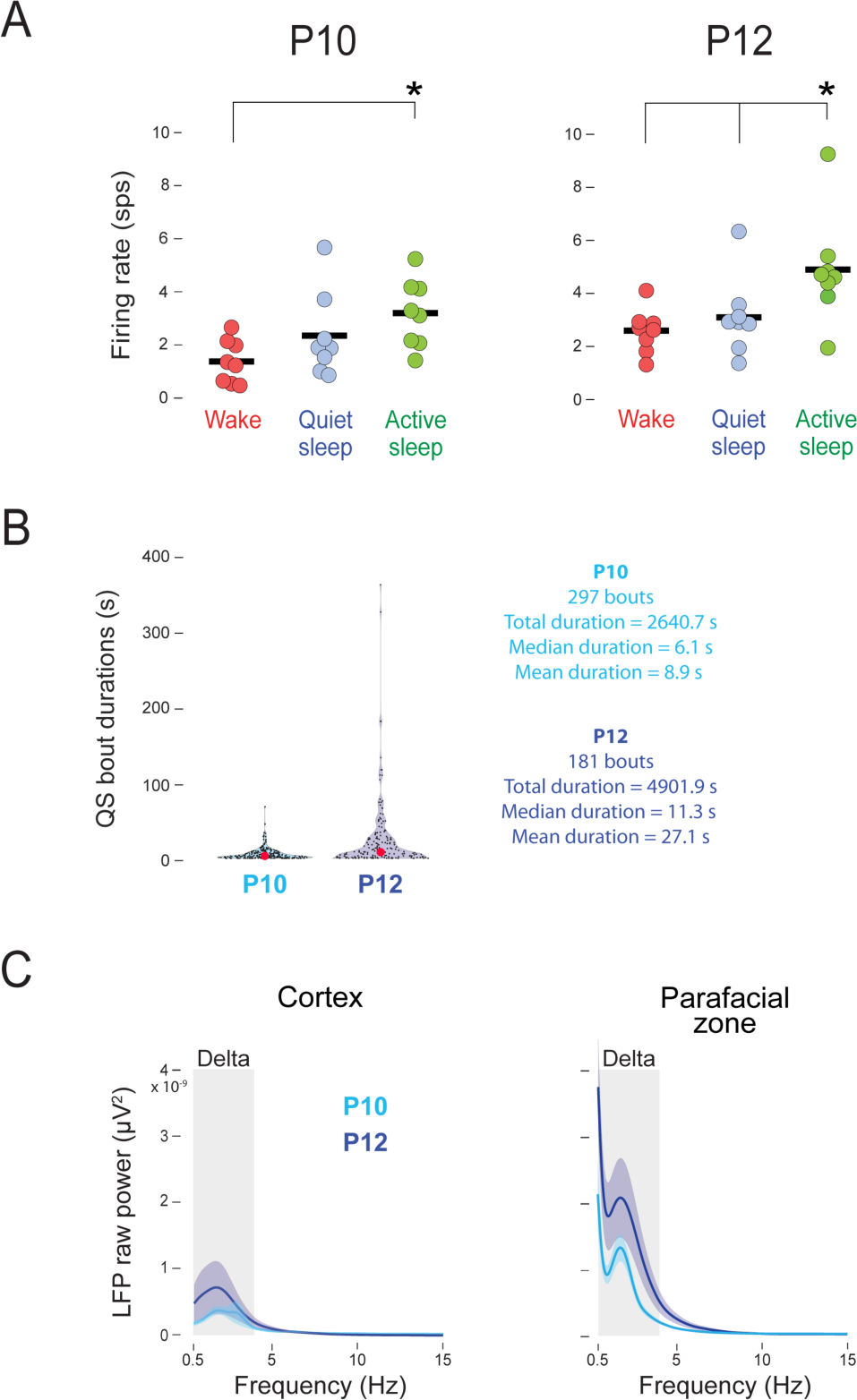
Developmental changes in PZ unit firing rates, QS bout durations, and raw delta power. Related to Figures 1, 3, and 6. (A) Mean firing rates (spikes/s) for each individual P10 (left) and P12 (right) rat during wake, quiet sleep, and active sleep. Individual data points are average values for pups. Asterisk denotes significant differences between groups. (B) Violin plots showing the durations of QS bouts across for all bouts in P10 (n = 8) and P12 (n = 8) rats. Red dots are median durations. (C) Mean (± SEM) raw power was computed in frontal cortex (left) and PZ (right) at P10 (n = 7) and P12 (n = 8). PZ delta was used to identify periods for analysis in both structures; however, the results were similar when cortical delta was used to define these periods (data not shown). Gray zones indicate delta frequencies (0.5-4 Hz).

**Figure S2.**
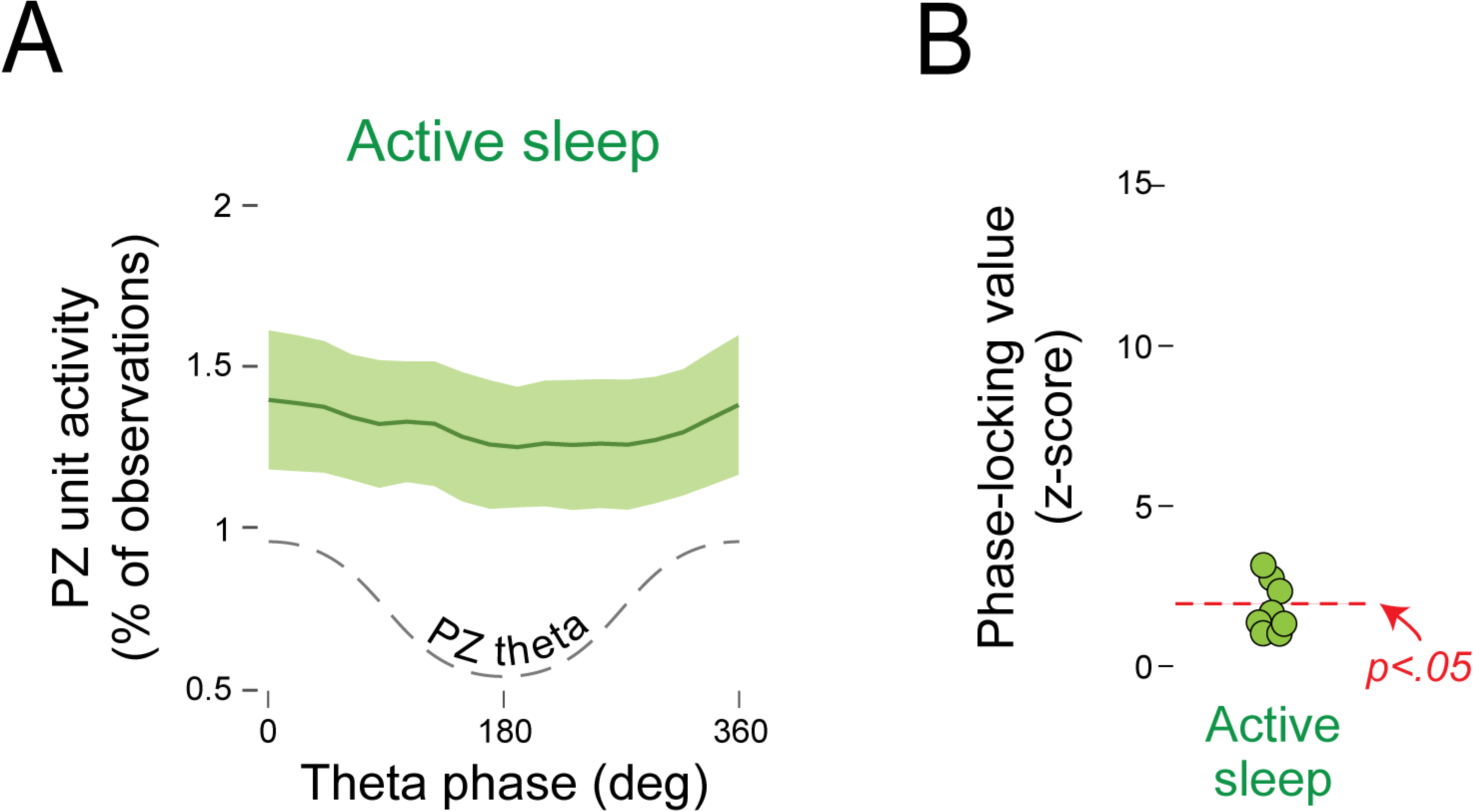
PZ units are not phase-locked with theta in P12 rats, Related to Figure 4. (A) Mean (± SEM) PZ unit activity in relation to theta phase in PZ during quiet sleep. Theta phase is illustrated with dashed lines. (B) Phase-locking values (z-scored) of PZ unit activity in relation to local theta phase during quiet sleep. Individual data points are average values for pups. Red dotted line indicates a significance level of p < .05 based on a z-score of 1.96.

**Figure S3.**
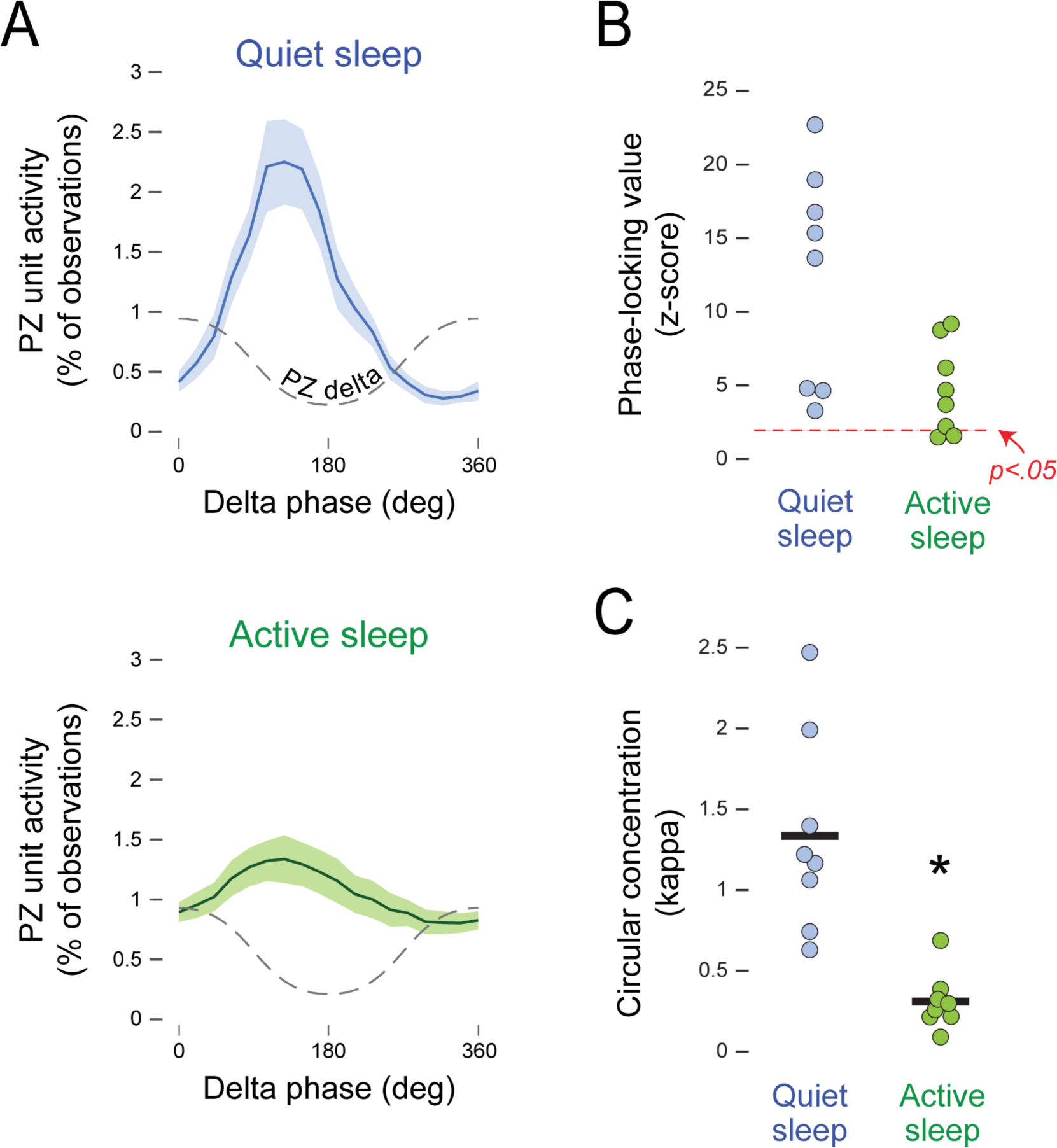
PZ units are phase-locked with delta during quiet sleep in P10 rats, Related to Figure 6. (A) Mean (± SEM) PZ unit activity in relation to delta phase in PZ during quiet sleep (top) and active sleep (bottom). Delta phase is illustrated with dashed lines. (B) Phase-locking values (z-scored) of PZ unit activity in relation to local delta phase during quiet and active sleep. Individual data points are average values for pups. Red dotted line indicates a significance level of p < .05 based on a z-score of 1.96. (C) Mean circular concentration coefficients (kappa) of PZ units in relation to local delta phase during quiet and active sleep. Individual data points are average values for pups. Asterisk denotes significant difference.

**Figure S4.**
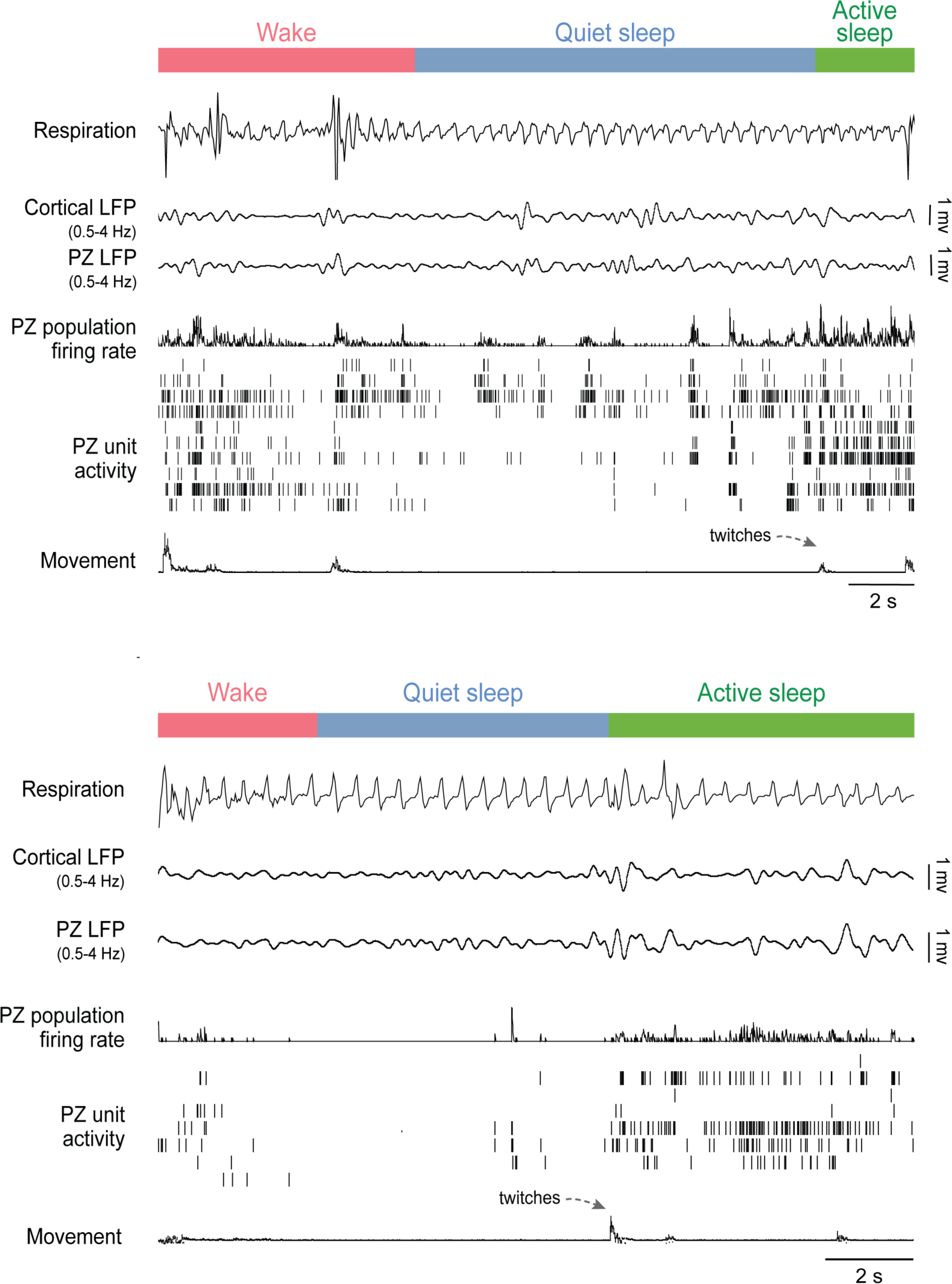
No evidence of delta-rhythmic PZ activity in P8 rats. Related to Figures 2 and 5. Representative recordings from two P8 rats cycling between wake (red), quiet sleep (blue), and active sleep (green). From top: respiration, cortical and PZ LFP, PZ population firing rate (i.e., the mean firing rate across all PZ single units), PZ unit activity, and limb movement.

**Figure S5.**
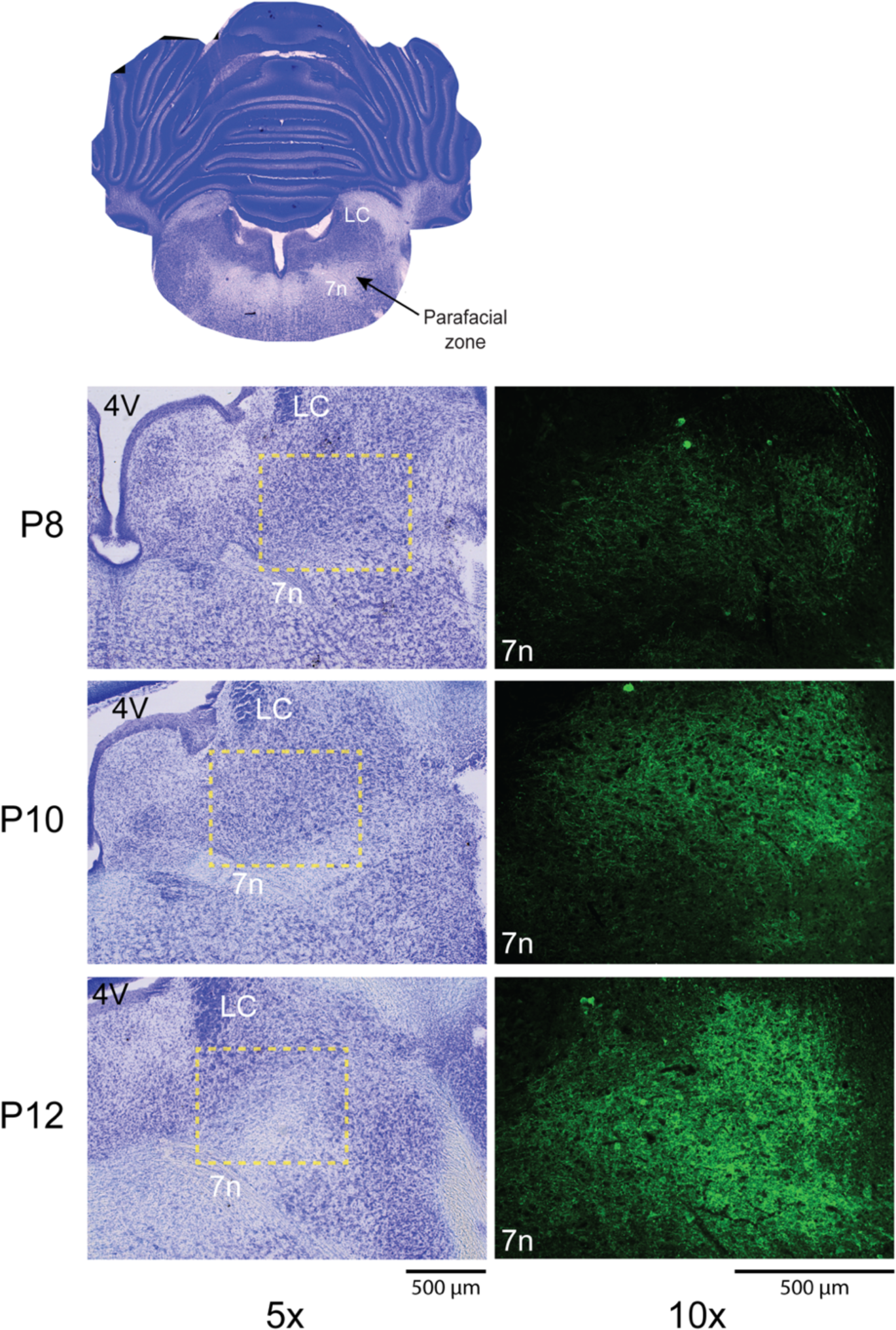
Increasing PV-expression in PZ between P8 and P12, Related to Figures 2 and 5. (A) Coronal section of brain of infant rat, showing the location of PZ immediately dorsal to the facial nerve (7n) and ventral to the locus coeruleus (LC). (B) Left column: Nissl-stained sections at 5x showing the region around PZ at P8 (top row), P10 (middle row), and P12 (bottom row). Dashed yellow boxes outline the regions shown at right: Right column: Representative sections at 10x showing PV-expressing terminals in PZ at the three ages. The exposure and gain were standardized across ages to enable age-related comparisons in expression. 4V, 4^th^ ventricle

**Figure S6.**
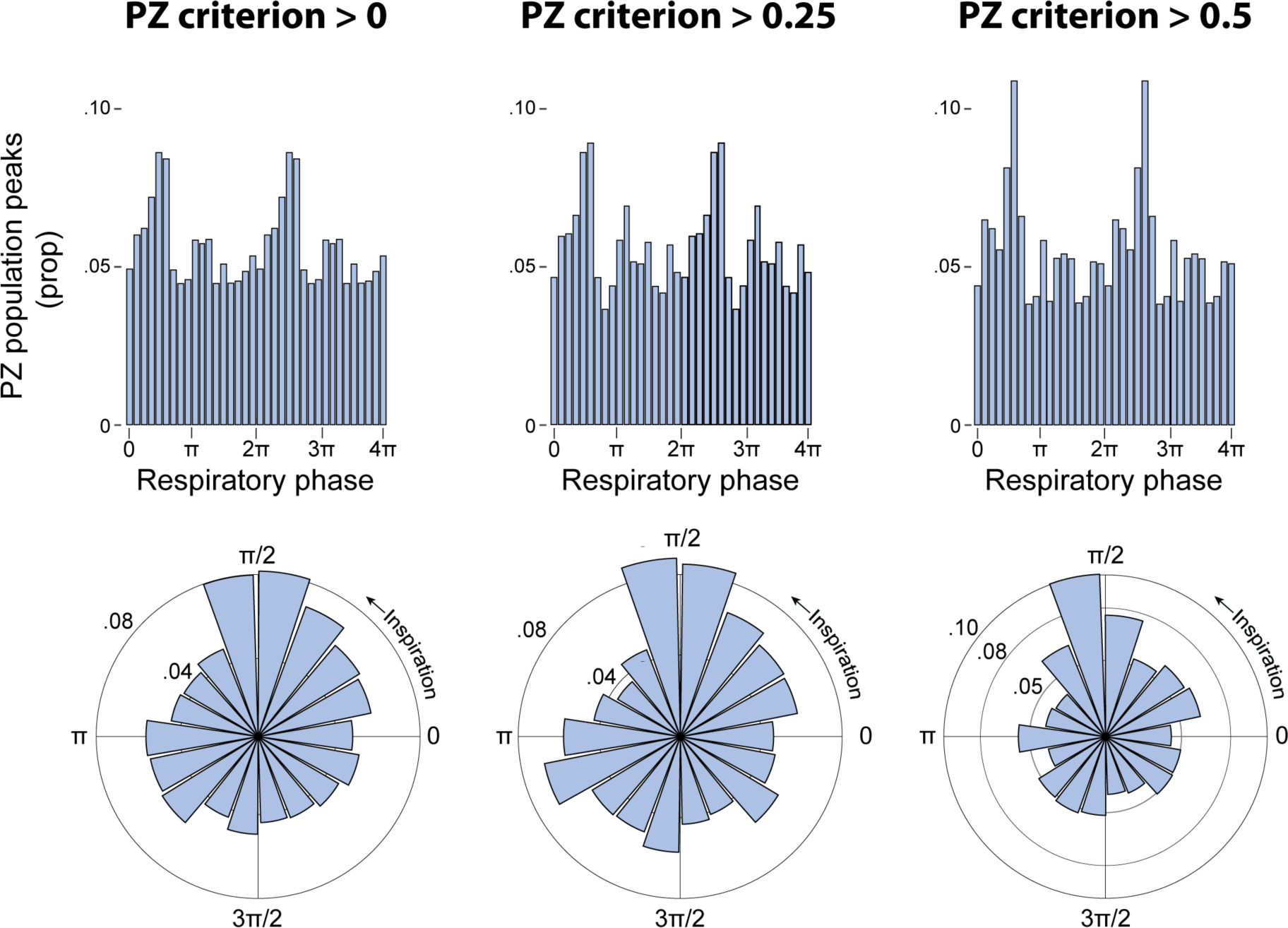
PZ population activity is phase-locked to respiration in P12 rats, Related to Figure 7. Top: Frequency distribution of mean proportion (prop) of PZ population peaks in relation to respiratory phase during quiet sleep. Note that the data are double-plotted. n = 8 pups. Bottom: Rose plots of the data above. Columns from left to right show results using increasingly stringent criteria for defining PZ bursting activity; the plots in the right-most column are the same as those in Figure 7D. The three distributions are significantly non-uniform (*Z*s ≥ 44.43, *p*s < .001).

**Figure S7.**
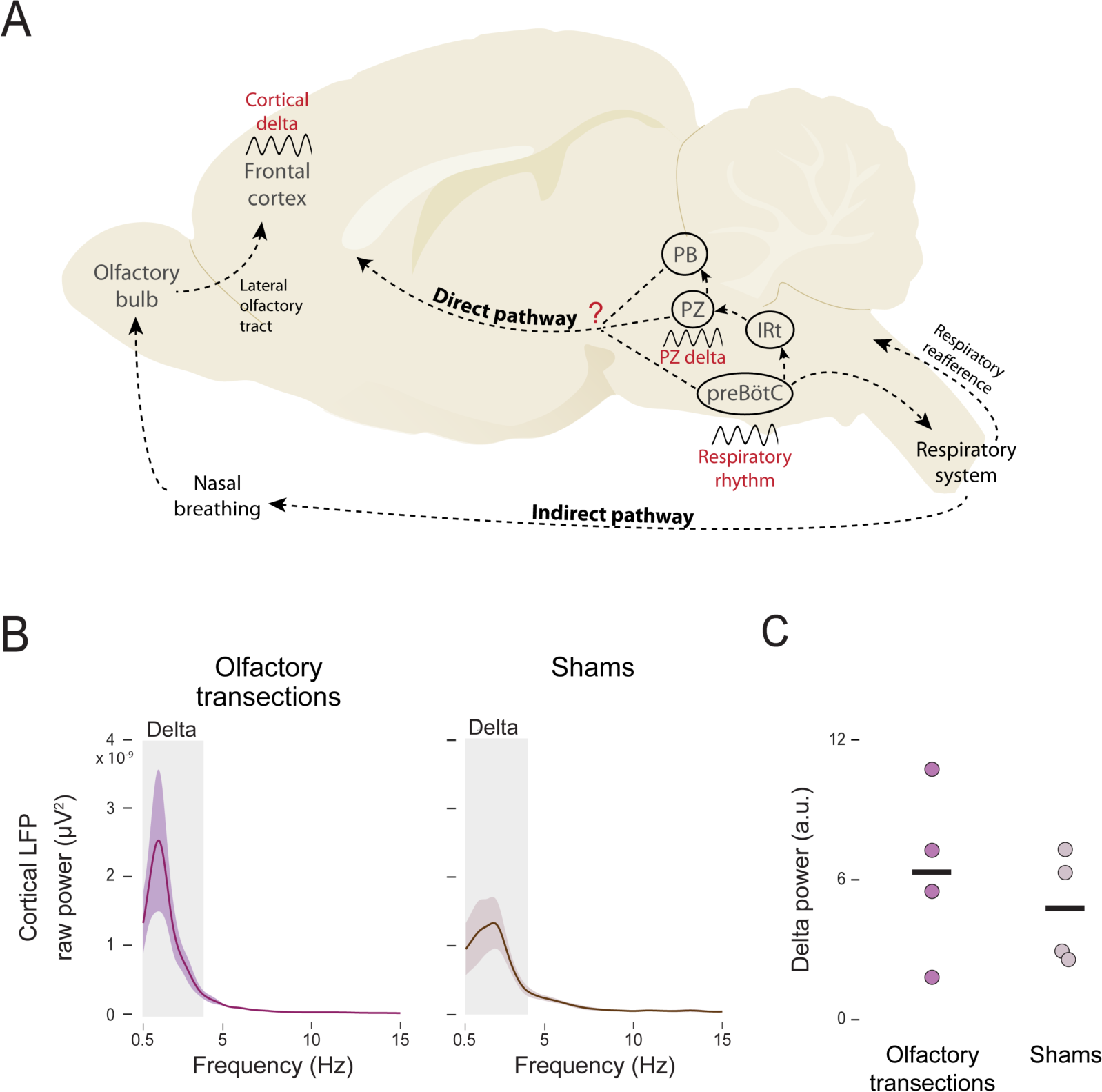
Disconnecting the main olfactory bulbs from the cortex at P12 does not diminish cortical delta power during quiet sleep, Related to Figure 7. (A) Possible mechanisms mediating the respiratory entrainment of PZ and cortical delta in infant rats. Breathing-entrained rhythmic spiking and delta in the parafacial zone (PZ) may arise from the inspiratory pacemaker in the preBötzinger complex (preBötC) via the intermediate reticular nucleus (IRt). PZ, preBötC, and the parabrachial nucleus (PB)—for example—have ascending projections that could directly influence how respiration entrains cortical delta. Also, in adults, it is thought that there is an indirect influence of respiration on cortical delta via nasal breathing and its effect on oscillatory activity in the olfactory bulbs. Finally, an influence of sensory feedback (reafference) from respiratory muscles on PZ activity is also possible. Adapted from Karalis and Sirota, 2022.^25^ (B) Mean (± SEM) raw cortical power in P12 rats after bilateral transection of the lateral olfactory tracts (left; n = 4) or sham transection (right; n = 4). Gray zones indicate delta frequencies (0.5-4 Hz). (C) Mean power (arbitrary units, a.u.) within delta frequencies for pups after olfactory or sham transection. Data points show power for individual pups.

## Notes

### Competing Interest Statement

The authors have declared no competing interest.

### Summary of Updates

This version reflects revisions following peer review. Changes were made primarily to the main text and to new figures. Also, some new data were added.

